# COVID-19 Vaccine Candidates: Prediction and Validation of 174 SARS-CoV-2 Epitopes

**DOI:** 10.1101/2020.03.20.000794

**Authors:** Marek Prachar, Sune Justesen, Daniel Bisgaard Steen-Jensen, Stephan Thorgrimsen, Erik Jurgons, Ole Winther, Frederik Otzen Bagger

**Author notes:** Correspondence to Frederik Otzen Bagger.

## Abstract

The recent outbreak of SARS-CoV-2 (2019-nCoV) virus has highlighted the need for fast and efficacious vaccine development. Stimulation of a proper immune response that leads to protection is highly dependent on presentation of epitopes to circulating T-cells via the HLA complex. SARS-CoV-2 is a large RNA virus and testing of all overlapping peptides *in vitro* to deconvolute an immune response is not feasible. Therefore HLA-binding prediction tools are often used to narrow down the number of peptides to test. We tested 19 epitope-HLA-binding prediction tools, and using an *in vitro* peptide MHC stability assay, we assessed 777 peptides that were predicted to be good binders across 11 MHC allotypes. In this investigation of potential SARS-CoV-2 epitopes we found that current prediction tools vary in performance when assessing binding stability, and they are highly dependent on the MHC allotype in question. Designing a COVID-19 vaccine where only a few epitope targets are included is therefore a very challenging task. Here, we present 174 SARS-CoV-2 epitopes with high prediction binding scores, validated to bind stably to 11 HLA allotypes. Our findings may contribute to the design of an efficacious vaccine against COVID-19.

## Introduction

2019-nCoV (SARS-CoV-2) was first reported in Wuhan, China, on 31 December 2019, following a series of unexplained pneumonia cases (WHO 2020a). Currently, the disease is intensifying with case reports over a continuously growing geographical area. WHO now gives the risk assessment ‘Very High’ on a global level and classifies the situation as pandemic (WHO 2020b, [c] 2020). Vaccine development is of high priority at this stage, and a number of public and private initiatives are focused on this task (Chen et al. 2020). Most, if not all, ongoing vaccine development efforts are focused on raising an immune response against the spike protein. However, the spike protein only makes up 1/8 of the COVID-19 genome, so this vaccine strategy may inadvertently miss a lot of potential immune reactivity. SARS-CoV-2 has a large proteome (F. Wu et al. 2020). Immune deconvolution to identify T cell epitopes will require initial filtering to assess which SARS-CoV-2-derived peptides are likely to bind a given HLA allotype and to be presented on the surface of infected cells from where it can activate passing T cells. The core binding groove of most MHC molecules can accommodate 9 amino acid residues, with some variation or suspected impact of flanking positions (Bassani-Sternberg et al. 2016; Rammensee 1995).

Several tools (a selection is presented in Table 1) have been developed that can predict the binding of peptides to HLA. Traditionally, these tools were trained using data from affinity assays (Harndahl et al. 2009), but more recently many of them also incorporate data from peptides identified by HLA ligandome analysis. Most tools rely on small neural networks (NN) or variations of position-specific weight matrices (PSSM), to calculate the probability of a peptide matching a consensus motif or model.

**Table 1.**
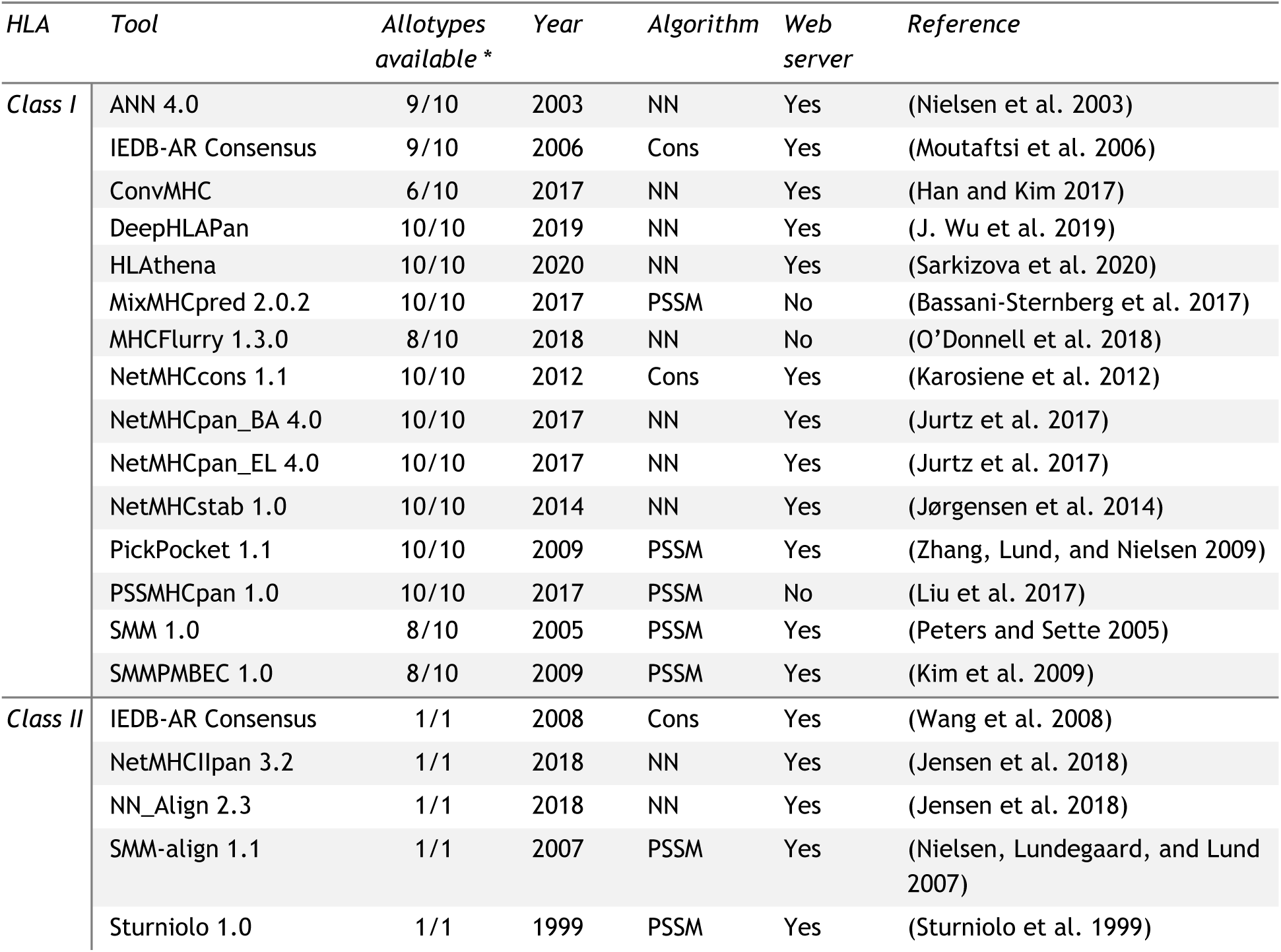
Current best-performing or novel HLA prediction tools (Mei et al. 2019). Webservers checked on 2 March 2020, NN: Neural network, Cons: Consensus, PSSM: Position specific scoring matrix. *availability as a fraction of allotypes included in study (10 HLA class I and 1 of HLA class II)

NetMHC tools (such as netMHC, netMHCII, netMHCpan, netMHCIIpan and others) have been under constant development and have consistently performed well throughout the last decade, (Peters, Nielsen, and Sette 2020; Mei et al. 2019; Saethang et al. 2012; Bhattacharya et al. 2017). Several tools are restricted in terms of which allotypes are available for prediction, in particular for MHC class II. This restriction is primarily determined by the availability of training data, for which the large public collection is currently the Immune Epitope Database (IEDB) (Sidney et al. 2006). Attempts to overcome this limitation have been made via sequence-to-sequence predictions, most notably for netMHCpan (Jurtz et al. 2017). A number of recent publications makes use of prediction tools to suggest vaccine candidate epitopes for SARS-CoV-2 (Fast, Altman, and Chen 2020; Grifoni et al. 2020; Abdelmageed et al. 2020).

To assess whether current peptide-HLA prediction tools could be suitable for identification of epitopes relevant in a vaccine against SARS-CoV-2, we tested predicted binders from the netMHC tools, using a new MHC:peptide complex stability assay NeoScreen^®^. We found that algorithmically predicting binding between epitopes from SARS-CoV-2 and HLA outputs many complexes that turned out to exhibit low stability. Such peptides are thus very unlikely to elicit an immune response against SARS-CoV-2 and are therefore unsuitable for vaccine development. To investigate if this finding was a result of the quality of available training data, we constructed a proof-of-concept prediction model, which we trained on 2,193 historic in-house stability data, and found that it outperforms other tools. Training data was primarily human cancer-derived or based on random sequences. SARS-CoV-2 peptides that we validated as binding or non-binding in this study are freely available for use to assist in vaccine design against COVID-19.

## Results

We set out to identify peptides with epitope potential in a future COVID-19 vaccine. We commenced by translating the reference sequence of SARS-CoV-2 (ACCESSION MN908947, VERSION MN908947.3) to protein-coding sequence and predicted potential epitopes in a sliding window of 9 using netMHC tools (netMHC/II and “-pan” versions, when allotype was not available). We identified the top 94 predicted peptides for 11 HLA allotypes (94 x 11 = 1034), and went further to validate the binding of these 94 peptides to each allotype in an *in vitro* MHC:peptide complex stability assay (NeoScreen^®^). We removed eight peptides that were synthetically introduced when translating the DNA sequence to protein sequence. Of the remaining 1,026 peptides we observed a high degree of overlap between different allotypes, resulting in 777 unique peptides. We found that 174 of the 777 unique peptides formed a stable peptide-HLA complex. Of these 174 peptides, 48 were previously measured and deposited in IEDB in relation to SARS and the remaining 126 peptides are novel. The full list of predicted binders (excluding synthetic peptides) can be found in the supplementary material (Supplementary Data S1).

In order to first assess potential variability across the stability measurements we made replicate measurements (*n*= 4) of 120 peptides on 8 HLA alleles. Each peptide was measured with urea in 4 different concentrations (0M, 2M, 4M, 6M), and we observed an average standard deviation between replicates of 0.10 with an average mean of 0.56. All remaining experiments were performed in duplicate for all concentrations.

To further address whether alternative prediction tools would have higher concordance with measured stability, we performed predictions for all tools listed in Table 1. Predictions for the 19 different tools were performed either through their web server or a stand-alone version, (see Materials and methods section for details). Furthermore, using in-house stability data, we developed PrdX 1.0, a prediction tool for a single allele HLA-A*02:01, where all other tools performed most poorly.

We assessed the false positive rate for each tool via Receiver Operating Characteristic (ROC) curves, and their Area under curve (AUC) for all allotypes that had more than 10 binders. The analysis revealed that ANN 4.0 achieved the highest score for allotype HLA-A*01:01 (AUC = 97,47; Figure 1A), closely followed by NetMHCcons 1.1, NetMHCpan_BA 4.0 and IEDB-AR Consensus. PrdX 1.0 scored highest for HLA-A*02:01 (AUC = 85.54; Figure 1B), NetMHCcons 1.1 scored highest for HLA-A*03:01 (AUC = 79.25; Figure 1C), and MHCflurry 1.3.0 performed best for HLA-B*40:01 (AUC = 91.06; Figure 1F). NetMHCstab 1.0 was the only tool that achieved the highest score for more than 1 allotype: HLA-A*11:01 and HLA-A*24:02 (AUC = 89.80; 86.03; Figure 1D, E, respectively). Out of the tools tested for HLA class II, IEDB-AR Consensus achieved the highest score for HLA-DRB1*04:01 (AUC = 81,31; Figure 1G). Table 2 provides all AUC values, and the result obtained for each allotype is marked in bold. Notably, in the case of HLA-A*02:01 we observed particularly poor performance among all tested tools despite the extensive amount of data available for this allotype.

**Table 2.** AUC values for ROC curves from Figure 1 for allotypes with more than 10 stable complexes. First 6 six columns contain HLA class I, last column contains HLA class II. Only four tools are tested for HLA class II. PrdX 1.0 is only available for allotype A*02:01. Highest value for each allotype is marked in bold.

| *Tool / allotype* | *A\*01:01* | *A\*02:01* | *A\*03:01* | *A\*11:01* | *A\*24:02* | *B\*40:01* | *DRB1\*04:01* |
| --- | --- | --- | --- | --- | --- | --- | --- |
| *ANN 4.0* | **97,47** | 70,3 | 77,06 | 83,81 | 82,96 | 89,76 | - |
| *IEDB-AR Consensus* | 97,06 | 69,44 | 77,36 | 87,05 | 83,89 | 90,03 | **81,31** |
| *ConvMHC* | 85,13 | 47,71 | 72,53 | 76,33 | 66,88 | 79,80 | - |
| *DeepHLAPan* | 91,82 | 62,90 | 52,49 | 80,84 | 69,87 | 80,95 | - |
| *HLAthena* | 89,74 | 65,41 | 66,54 | 87,41 | 76,48 | 83,08 | - |
| *MixMHCpred 2.0.2* | 92,54 | 70,04 | 75,29 | 80,82 | 78,68 | 76,29 | - |
| *MHCFlurry 1.3.0* | 94,48 | 66,88 | 75,52 | 88,46 | 76,24 | **91,06** | - |
| *NetMHCcons 1.1* | 97,42 | 65,93 | **79,25** | 86,21 | 79,52 | 90,25 | - |
| *NetMHCpan_BA 4.0* | 97,15 | 65,84 | 76,32 | 86,76 | 85,40 | 88,79 | - |
| *NetMHCpan_EL 4.0* | 93,40 | 75,89 | 75,84 | 80,11 | 78,36 | 84,27 | - |
| *NetMHCstab 1.0* | 89,15 | 76,75 | 77,98 | **89,80** | **86,03** | 86,13 | - |
| *PickPocket 1.1* | 88,65 | 57,32 | 65,53 | 75,03 | 80,93 | 88,44 | - |
| *PrdX 1.0* | - | **85,54** | - | - | - | - | - |
| *PSSMHCpan1.0* | 90,33 | 67,97 | 75,75 | 76,62 | 82,12 | 77,94 | - |
| *SMM 1.0* | 95,25 | 60,09 | 75,15 | 87,26 | 80,26 | 88,82 | - |
| *SMMPMBEC 1.0* | 92,13 | 60,48 | 75,36 | 87,26 | 80,13 | 86,34 | - |
| *NetMHCIIpan 3.2* | - | - | - | - | - | - | 76,63 |
| *NN_Align 2.3* | - | - | - | - | - | - | 78,14 |
| *SMM-align 1.1* | - | - | - | - | - | - | 74,42 |
| *Sturniolo 1.0* | - | - | - | - | - | - | 75,19 |

**Figure 1.**
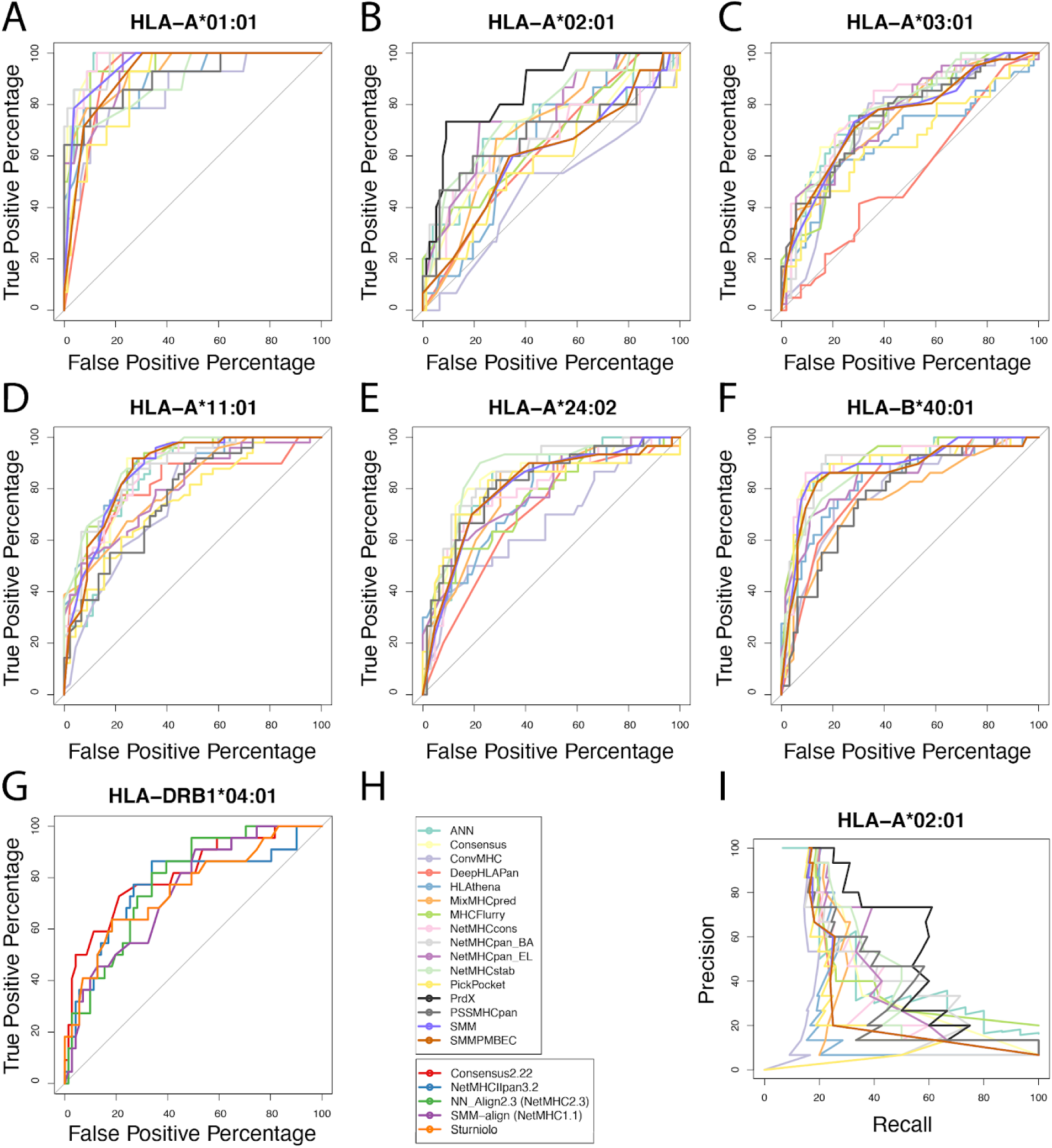
ROC curves for each allotype that bound more than 10 peptides stably (subplots A, B, C, D, E, F, G), H) tools used in the benchmark, I) precision-recall curves for HLA-A*02:01. Corresponding area under curve (AUC) values are listed in Table 2.

To assess the correlation between the predicted and measured peptide-HLA complexes, Spearman correlation was calculated for all allotypes. This revealed significant inconsistencies in performance depending on the predicted allotype. PSSMHCpan 1.0 displayed the highest consistency, taking into account its coverage (Table 1), but the correlation median scored lower than other tools such as IEDB-AR Consensus, MixMHCpred 2.0.2, NetMHCpan_EL 4.0 or PrdX 1.0. The results of the Spearman correlations are summarised in Figure 2.

**Figure 2.**
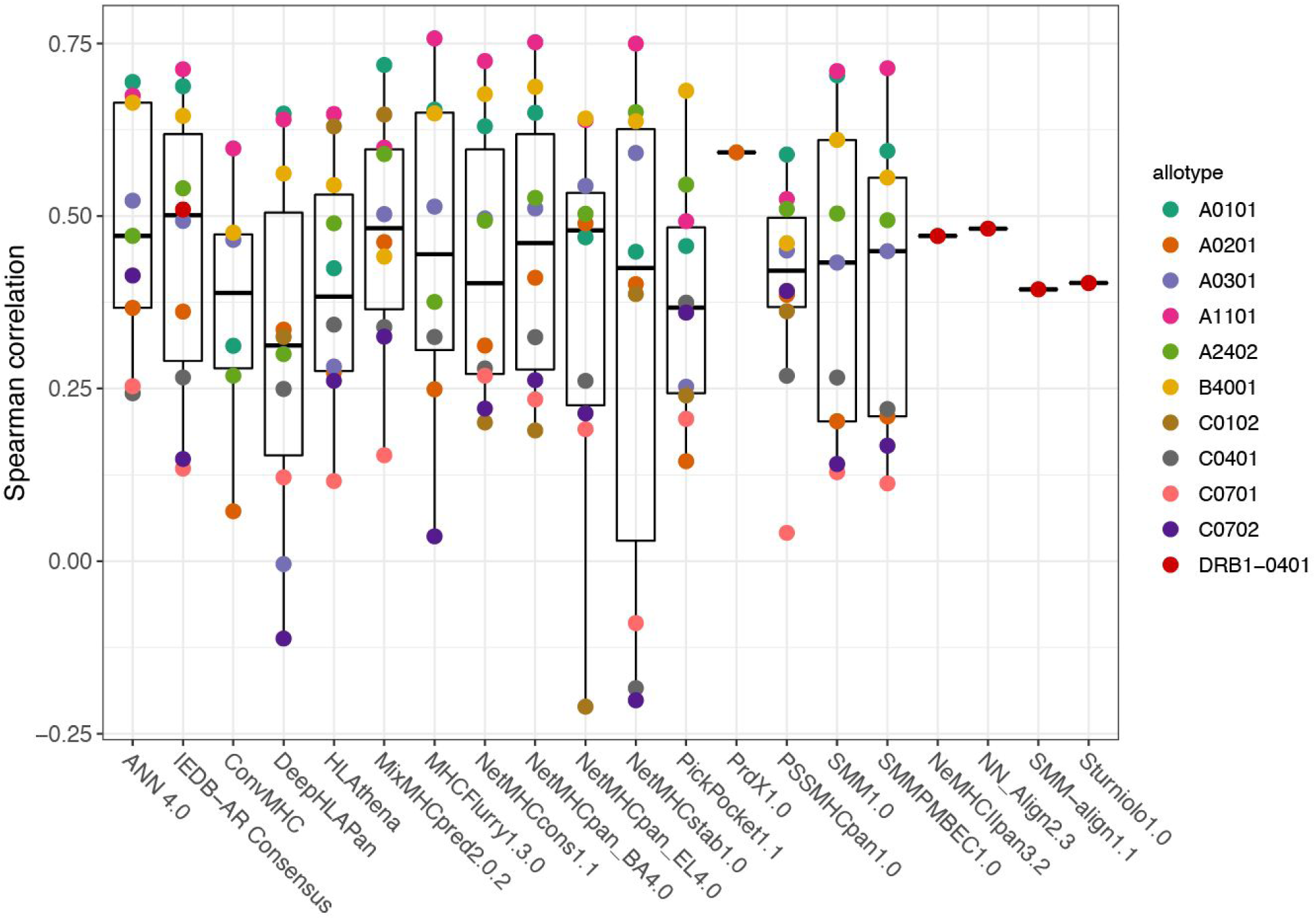
Plot of Spearman correlation between predicted values and results of Neoscreen^®^ stability assay for each available allotype. Each colour represents an individual allotype. Whiskers accord for 1.5 distance between the median and quartile hinges. Data points beyond the end of the whiskers are outliers.

## Discussion

Here we benchmark a number of tools to identify epitopes for SARS-CoV-2 virus, and validate via stability assay the binding of candidate epitopes to 10 allotypes of HLA class I and one allotype of HLA class II. We find that the false positive rate is high for all tested tools when testing for predicted HLA-binding peptides from SARS-CoV-2 virus. This creates a challenge for vaccine development efforts, especially for the design of epitope vaccines, where only a limited number of epitopes may be included. Furthermore, it highlights the risk for failed vaccine design (for any pathogen or disease) if predicted HLA-binding protein regions in reality do not bind and allow immune presentation and response.

We observed, remarkably, that all tools tested performed poorly for HLA-A*02:01, which is the allotype with most training data available (Kim et al. 2014). Based on our observations we hypothesise that publicly available training data is not of high enough quality. This is supported by the fact that AUC and Spearman correlations indicate that performance seems to correlate with the allotypes and not the tools; thus suggesting that either the training data or the difficulty of modelling the allotype is responsible for poor predictions. To test the hypothesis that training data is limiting for tool performance, we trained a vanilla NN on only 2,193 historic in-house stability measurements, and found that our model outperformed all tested prediction tools in this setting. This observation could also be explained by more similar data distributions between test and training data for PrdX 1.0.

We identified 174 potential SARS-CoV-2 vaccine candidate peptides, out of which 48 have been previously deposited in IEDB following various studies (Qu et al. 2019; Blicher et al. 2005; Sylvester-Hvid et al. 2004; Harndahl et al. 2006; Sidney et al. 2006; Ishizuka et al. 2009). The majority of the previously deposited peptides were measured in one or multiple affinity assays and reached low Kd (<50 nM) values, indicating strong affinity. Additionally, 9 of these peptides were previously measured in another stability assay and were recognised as stable binders (Rasmussen et al. 2014), independently confirming our approach and measurements. Of the 48 peptides deposited in IEDB, two (FLLPSLATV https://www.iedb.org/epitope/16743, FLNRFTTTL https://www.iedb.org/epitope/16786) have previously been assessed for their ability to provoke T cell responses, with negative T cell responses for both.

To further assess the potential of the SARS-CoV-2 epitopes we explored their overlap with T cell-confirmed epitopes from SARS that are deposited in the IEDB. We identified 4 peptides within our candidates that are represented within these confirmed epitopes as partial motifs.

In conclusion, we make freely available the identities of 174 COVID-19 epitopes that we have predicted and validated *in vitro* to be HLA-binding. We hope that this contribution will aid the development of a vaccine against SARS-CoV-2. We performed a benchmark analysis of 20 tools on 777 peptides that were predicted by state-of-the-art prediction tools from netMHC to be binders, and revealed high false positive rates for all benchmarked tools. We observed improved performance after training our own prediction tool PrdX 1.0 on allotype A*02:01 using in-house generated stability data. Our findings suggest that the performance of current state-of-the-art epitope prediction tools are impacted by the varying quality of publicly available data.

## Data availability

All epitopes are available at the vendor webpage (www.immunitrack.com) and in supplemental materials (Data S2.).

## Materials and methods

Fifteen prediction tools tested on a relevant dataset of peptides from the SARS-CoV-2 genome (assembly MN908947.3). The genome sequence was downloaded from the NCBI database (https://www.ncbi.nlm.nih.gov/nuccore/MN908947.3) (F. Wu et al. 2020). Using NetMHC tools we predicted the top 94 peptides for HLA-A*01:01, HLA-A*02:01, HLA-A*03:01, HLA-A*24:02, HLA-B*40:01, HLA-C*04:01, HLA-C*07:01, HLA-C*07:02 (netMHC 4.0), HLA-C*01:02 (netMHCpan 4.0) and HLA-DRB1*04:01 (netMHCII 2.3). Subsequently, the peptides were analysed for binding stability to the respective HLA allotype. Taking into account the cross-reactivity between the two allotypes, peptides predicted to bind HLA-A*03:01 were also measured on HLA-A*11:01.

Peptides were synthesised using standard Fmoc solid-phase synthesis on a modified cellulose support as solid support according to the SPOT synthesis protocol, starting with the acid labile Ramage linker.

After synthesis, peptides were cleaved off the membranes using 95% trifluoroacetic acid (TFA),3% triisopropylsilane (TIS) & 2% H2O.Peptides were then precipitated with diethylether and washed with methyl-tert-butylether.

Peptides were subsequently dissolved in a proprietary mixture and dried under vacuum using a speed vac. Finally, 5% of all peptides were analysed by MALDI-TOF to confirm correct molecular weight. The anticipated yield per spot was 50 μg.

### NeoScreen^®^ assay

The NeoScreen^®^ stability assay utilises urea denaturation to assess peptide:MHC complex stability. Briefly, peptides were dissolved in 200 µl DMSO with 1 mM β-mercaptoethanol and subsequently diluted into an assay buffer in 96 well plates at a final concentration of 2 µM. Positions A1 and H12 were reserved for a mixture of reference peptides with known stable binding to the MHC of interest. MHC I was diluted into an assay buffer with beta 2 microglobulin and added at a 1:1 ratio to diluted peptides. For MHC II, the urea-denatured alpha and beta chains were diluted into an assay buffer and added at a 1:1 ratio to diluted peptides. The concentration of MHC depended on the actual chain but final concentrations were in the range of 2-10 nM (hence peptide was added in excess). Upon folding, peptide:MHC complexes were transferred to 384 well plates where they were challenged with 4 different urea concentrations. Following the period of urea-induced stress the plates were developed in a conventional ELISA as described previously (Justesen et al. 2009; Sylvester-Hvid et al. 2002). The ABS450 nm signals from the 4 different wells were averaged and normalised to the included reference to the included reference peptides in wells A1 and H12.

### Benchmarking of tools

Table 1 provides a summary of tools tested in this benchmark analysis. It features the year of their development, the algorithm used, web server availability and a reference. Most of the tested tools are available at the IEDB Analysis Resource web page (http://tools.iedb.org/main/) and were run through their web interface (http://tools.iedb.org/mhci/ or http://tools.iedb.org/mhcii/). MixMHCpred, MHCflurry and PSSMHCpan were downloaded from their respective github pages (https://github.com/GfellerLab/MixMHCpred,https://github.com/openvax/mhcflurry, https://github.com/BGI2016/PSSMHCpan, respectively). ConvMHC, DeepHLApan and HLAthena were used from their privately hosted web servers (http://jumong.kaist.ac.kr:8080/convmhc, http://biopharm.zju.edu.cn/deephlapan/, http://hlathena.tools/, respectively).

All tested peptides were subjected to in silico predictions (with each prediction tool) regarding their available allotype. Predictions were compared against measured stability determinations obtained through the NeoScreen^®^ assay. Measurements were normalised to an allotype-specific reference peptide (stability = 100). The list of reference peptides used is available in supplementary material (Table S1.). The threshold for a stable binder was set to 60. Predictions were subsequently evaluated according to commonly-used metrics such as the Receiver Operating Characteristic (ROC) and its Area Under Curve (AUC) to visualise the relationship between sensitivity and specificity. Spearman correlation was also used to compare the ranked correlation of predicted and measured data.

### PrdX

To assess the performance of predictors trained on stability data we used PyTorch (Paszke et al. 2017) to train a fully connected, feed-forward neural network with 64 and 32 hidden units on historic in-house stability data from allotype A*02:01. This data contains a mixture of human cancer-related stability measurements and measurements made on synthetic random peptides. We used BLOSUM62 matrix for encoding, simple network architecture, train-test split and early stopping for training.

