## Supplementary Materials for "COVID-19 Vaccine Candidates: Prediction and Validation of 174 SARS-CoV-2 Epitopes"

This PDF file includes:

Materials and Methods
Table S1
Captions for Data S1-S2

### Materials and Methods

#### Analysis of predictions

To produce the ROC graphs and to calculate their AUC we used RStudio Version 1.2.5033, R version 3.6.2, pROC 1.16.1 package. The same software was used to calculate and plot Spearman correlations, using the ggplot2 3.2.1 package.

#### PrdX

The software was written in Python Version 3.7.3, NN architecture was built using PyTorch Version 1.3.0.

Table S1. List of reference peptides used for their corresponding allotype .

| Allotype | Sequence | Reference |
|---|---|---|
| *A*0101* | VTEHDTLLY | http://www.iedb.org/epitope/71290 |
| *A*0201* | VLDFAPPGA | Wilms tumor antigen 1 |
| *A*0301* | AVAHKVHLMYK | https://www.iedb.org/epitope/419554 |
| *A*1101* | AVFDRKSDAK | https://www.iedb.org/epitope/5316 |
| *A*2402* | AYAQKIFKIL | https://www.iedb.org/epitope/5731 |
| *B*4001* | REDQWCGSL | https://www.iedb.org/epitope/53476 |
| *C*0102* | QYDPVAALF | http://www.iedb.org/epitope/52886 |
| *C*0401* | QYDPVAALF | http://www.iedb.org/epitope/52886 |
| *C*0701* | YLHARLREL | Identified by previous screening |
| *C*0702* | NYFNRMFHF | Identified by previous screening |
| *DRB1*0401* | AKFVAAWTLKAAA | https://www.iedb.org/epitope/2192 |

Data S1. COVID19-Intavis-Immunitrack-dataset

https://www.immunitrack.com/wp/wp-content/uploads/Covid19-Intavis-Immunitrack-datasetV2.xlsx

Excel file containing results of the NeoScreen® assay.

Data S2. Full list of predicted and measured peptides used in the benchmark

https://doi.org/10.5281/zenodo.3739310

Public datasets of all peptides that were predicted and subsequently measured. List of peptide sequences for each respective allotype. All predictions of the benchmarked tools, not all tested tools use the same measure.
